# CARP2 regulates the Golgi dynamics upon EGF stimulation

**DOI:** 10.1101/2022.04.05.487156

**Authors:** Rahul Sharma, Srinivasa M. Srinivasula

**Author notes:** Correspondence to Srinivasa M. Srinivasula.

## Abstract

Golgi apparatus regulate diverse cellular functions like protein sorting, vesicular trafficking, secretion, protein modifications like glycosylation etc. In mammalian cells though, Golgi apparatus appear as ribbon architecture, individual stacks laterally linked to each other by tubular structure, it’s architecture changes dynamically to cater to the needs of the cell under physiological and stress conditions. Loss of Golgi integrity is reported to be associated with pathological conditions like cancer and neurodegeneration. Very little is known of molecular regulators of Golgi dynamics. Here, we demonstrate that CARP2 (Caspase −8 and −10 associated RING containing protein 2), an endosomal ubiquitin ligase and a known regulator of cell migration, modulates the Golgi structure. Stimulation with EGF (Epidermal growth factor) modestly increased CARP2 protein levels. CARP2 exogenous expression or EGF treatment resulted in dispersal of the Golgi apparatus. Conversely, CARP2 deletion suppressed EGF induced Golgi dispersal. CARP2 variants that are defective in their endosomal-association or E3 ligase activity were unable to exhibit Golgi dispersal, indicating importance of both the endosomal localization and the E3 activity for this function. Importantly, we provide evidence that in EGF stimulated cells CARP2 mechanistically functions by targeting one of the Golgi structural proteins, Golgin45 for ubiquitination and degradation. Taken together, our findings unravel the existence of crosstalk between endosomal ubiquitin signaling and the Golgi dynamics.

**Significance:** The Golgi is an organelle that exists in mammals in ribbon form - individual stacks laterally linked with each other - is central to protein and lipid modifications, trafficking and secretion. The Golgi architecture is changed dynamically to cater to the physiological needs of the cells (eg: cell division, migration). Dysfunctional or altered Golgi is reported under pathological conditions like cancer, neurodegenerative diseases etc. This study unravels a complex signaling between endosomal ubiquitin ligase, CARP2 and one of the Golgi structural proteins, Golgin45. Here, we delineate CARP2-Golgin45 signaling as a fundamental mechanism that regulates Golgi dynamics underlying in EGF-stimulated cell migration.

## Introduction

Dynamic reorganization of organelle, defined by their cellular localization, number, shape, and size is critical for spatio-temporal regulation of diverse cellular processes. These dynamics are in turn precisely coordinated by complex networks of molecular machinery^1, 2^. The dynamic nature of organelle like mitochondria, endoplasmic reticulum, and lysosomes, under physiological or pathological conditions, is relatively more investigated than the Golgi apparatus, which is the central hub for protein and lipid trafficking and secretion.

In mammalian cells, Golgi apparatus exhibit a high degree of complexity with a ribbon-like structure, mostly reported to be restricted to the perinuclear or pericentriolar region. Golgi ribbon consists of multiple stacks of a tightly packed flat-disc-shaped membranous cisterna, fused laterally by tubular structure or proteinaceous matrix^3–5^. The stack consists of 5-7 polarized cisternae, divided into distinct regions like cis, medial, and trans-Golgi. The cis-Golgi region is located towards the ER and receives output from ER. The cargo destined to intracellular or extracellular locations exit from the trans-Golgi region^6^. All the zones are involved in modification and sorting of the cargo. Recent findings establish that architecture of the Golgi apparatus switches between ribbon to dispersed/mini-stacks structures, to cater the needs of the cell (viz. cell division and migration)^7–9^. Moreover, this remodelling of the Golgi occurs under physiological conditions in a cell type-specific manner. For example, in neurons Golgi apparatus are present as outposts near dendritic outgrowth besides the perinuclear region. Golgi outposts are thought to regulate the dynamics of microtubules and the delivery of the cargo locally, whereas in muscle cells interspaced Golgi mini stacks act as microtubule nucleation sites in the myofibril networks^8, 10^.

Under pathological conditions like Alzheimer’s disease, Huntington’s disease, and Parkinson’s disease, disruption of the Golgi is also reported in cells of hippocampal and cortex tissues of the brain^11^. In the transgenic mouse model of AD, accumulation of beta-amyloid plaques induced Golgi fragmentation through downregulation of Golgi structure proteins GRASP65 and 55^12^. A number of known Golgi structural proteins viz GM130, Golgin45, Golgin97, Giantin, GRASP65 localize to distinct regions of the Golgi and regulate its dynamics primarily by acting as tethering molecules^7, 13–15^. The recruitment and trafficking of these Golgi structural proteins are regulated by GTPases like Rabs, Arf, and ARL1^13^. Besides these molecules, the cytoskeleton also reported to play a crucial role in vesicle trafficking as well as in maintenance of the Golgi architecture. Treatment with nocodazole (a microtubule depolymerization inhibitor) is linked with scattering of the Golgi apparatus throughout cytosol. On the contrary, treatment with cytochalasin D, (an actin inhibitor) is reported to make the Golgi structure more compact than extended ribbon^13, 16^. Moreover, signaling factors like EGF (Epidermal growth factor), histamine, cytosolic calcium levels, ER stress etc. are reported to affect the Golgi architecture^17, 18^. Nonetheless knowledge of molecular mechanism(s) that regulate Golgi organization under various physiological and pathological conditions is very limited.

Here, we report that an endosomal-associated ubiquitin ligase, CARP2(also known as RFFL) is a novel regulator of Golgi apparatus in mammalian cells. In EGF (Epidermal growth factor) stimulated cells, CARP2 contributes to the Golgi reorganization by targeting one of the Golgi structural proteins, Golgin45 for ubiquitination and degradation. We demonstrate that Golgin45 levels are reduced upon EGF stimulation in wildtype, but not in CARP2 KO cells. CARP2 ligase activity and endosomal association is essential for this function.

## Results

### EGF induced CARP2 upregulation leads to Golgi dispersal

Stimulation with hEGF (Epidermal growth factor) results in the activation of ERK/MEK (Extracellular-signal-regulated kinase / Mitogen-activated protein kinase) pathway and remodeling of the Golgi architecture^19^. Moreover, we have previously reported that activation of ERK/MEK leads to upregulation of CARP2 transcription^19, 20^. Hence, we hypothesized that CARP2 regulates EGF-dependent Golgi remodeling. To investigate this, first we treated A549 lung carcinoma cells (wildtype) with human EGF for different time points and evaluated endogenous CARP2 protein levels by western blotting (Figure 1A). As hypothesized, treatment with EGF resulted in enhanced CARP2 protein levels within 2 hours. As expected, no CARP2 protein was detected even after EGF treatment in A549 cells in which CARP2 expression was knocked out using CRSIPR/Cas9 strategy (CARP2 KO cells).

**Figure 1.**
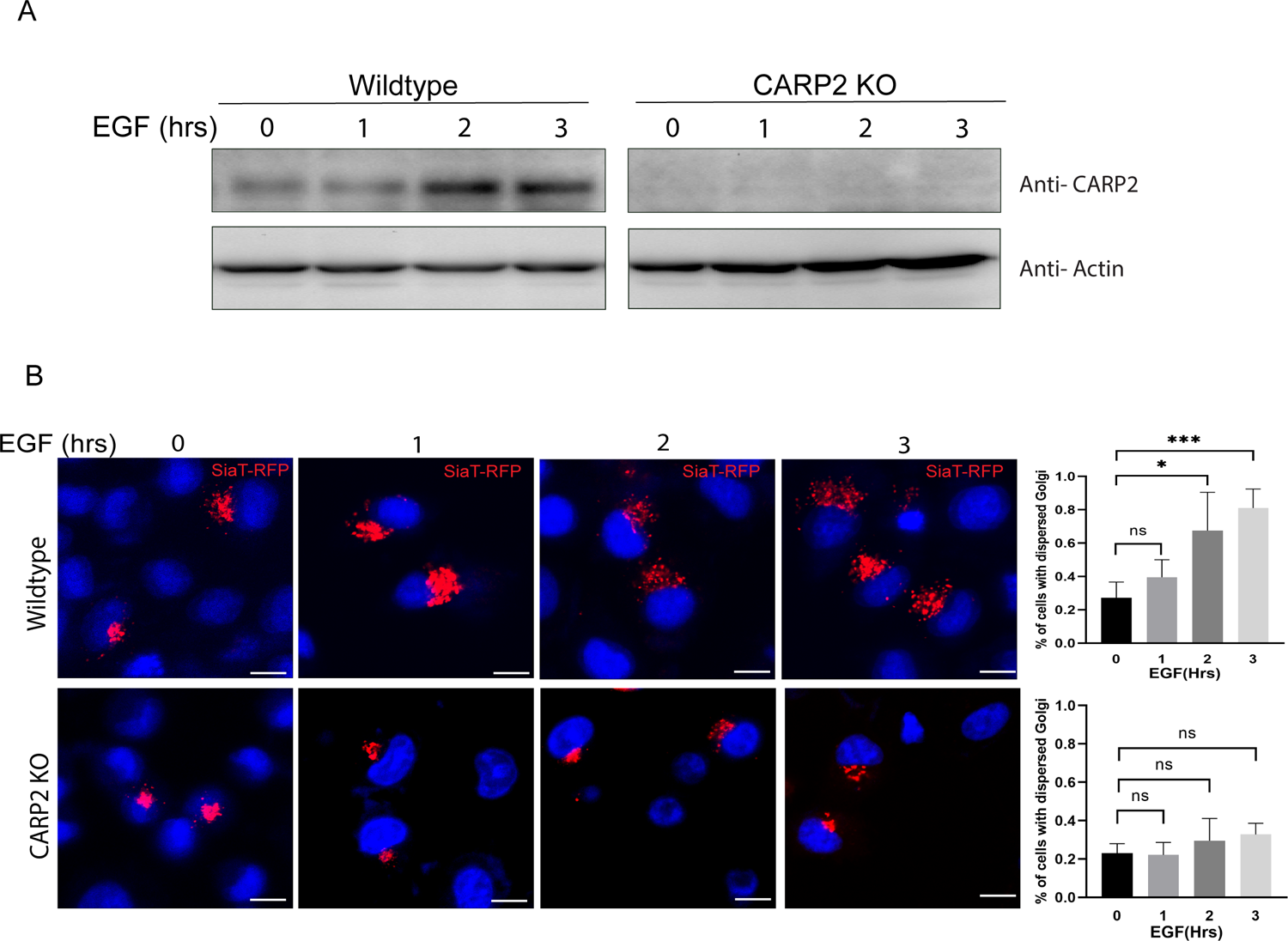
EGF induced CARP2 upregulation leads to Golgi dispersal. **A.** EGF treatment was given to A549 wildtype or CARP2 KO cells for indicated time periods and extracts were probed anti-CARP2 and anti-actin antibody. Representative blot is shown from three independent experiments. **B.** A549 or CARP2 KO cells transiently expressing SiaT-RFP were treated with EGF and imaged live at different time points. Nucleus was stained with a Hoechst stain (upper panel). Scale 10µm. Quantification of Golgi dispersal was shown in the lower panel. Statistical analysis was computed by ANOVA, ***,*P<0.001. Error bars represent mean ± SEM from three independent experiments. At least 50 cells were imaged for each biological replicate

Further, we evaluated the role of CARP2 in EGF induced Golgi dynamics. For this experiment, we employed SiaT-RFP to monitor Golgi dynamics. SiaT is a sialyltransferase enzyme known to localize to trans-Golgi region, and glycosylate numerous proteins within the Golgi. SiaT-RFP is a construct consisting of human Golgi localization sequence (nucleotide sequence corresponding to 1-110 amino acids) cloned in frame with RFP. Usage of this construct as a marker to monitor Golgi dynamics in cells has been well established^21^.

The distribution of SiaT-RFP, was limited mostly to the perinuclear region in both CARP2 wildtype and CARP2 KO cells without any treatment, like GM130 immunostaining (Figure S3). Consistent with an earlier report, EGF stimulation led to dispersal of SiaT-RFP in wildtype cells, indicating remodeling of the Golgi apparatus. However, this dispersal was substantially reduced in CARP2 knockout cells suggesting the important role of CARP2 in EGF dependent Golgi remodeling (Figure 1B).

### Exogenous CARP2 expression alone regulates the Golgi architecture

Since EGF stimulation increases endogenous CARP2 protein levels and CARP2 is required for Golgi dispersal in EGF treated cells, we wanted to test a possibility that an increase in the protein levels of CARP2 could affect the Golgi dynamics even without EGF stimulation. To investigate this hypothesis, we have evaluated the Golgi architecture in A549 cells stably expressing untagged CARP2 by immunostaining for GM130, a well-known Golgi marker. We noted that mere exogenous-expression of CARP2 resulted in dispersal of the GM130-positive structures whereas in vector control cells the Golgi stain appeared as compact. Quantification of the Golgi dispersal revealed that more than 80% of CARP2 expressing cells exhibited dispersed Golgi phenotype in comparison to 20% of the control cells (Figure 2A & B, S3). Staining of these cells with anti-CARP2 antibody showed localization to mostly intracellular endocytic vesicles, a pattern consistent with earlier published reports (Figure S1A). Results from immunoblots from the extracts of cells expressing either vector alone or CARP2 probed with anti CARP2 anti body are shown in figure S1A.

**Figure 2.**
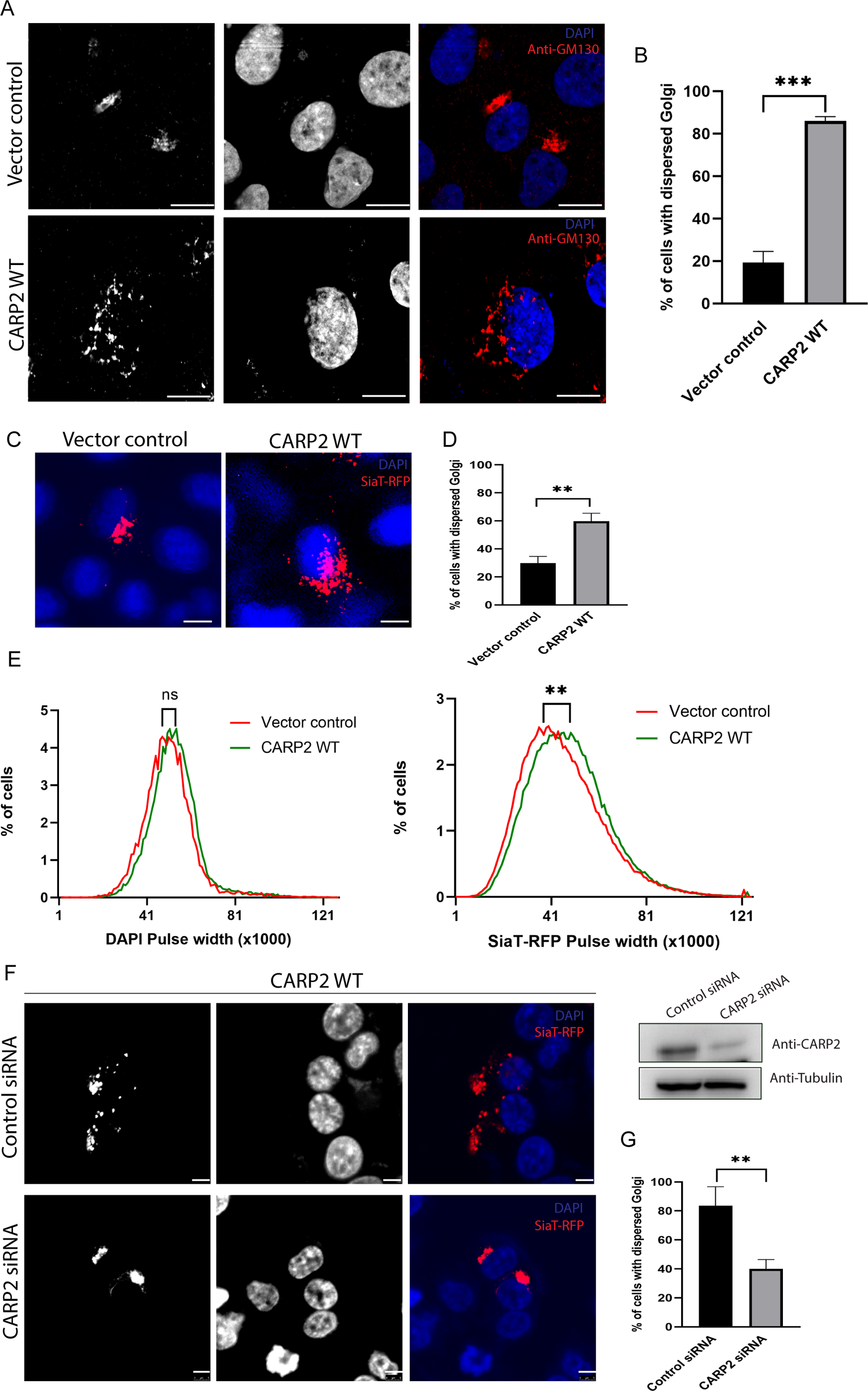
Expression of an endosomal-associated ubiquitin ligase CARP2 results in the dispersal of Golgi structure. **A.** A549 cells stably expressing either vector control or CARP2 without any tag were cells immunostained with GM130 antibody (red). Nucleus was stained with DAPI (nucleus). Scale 10µm. **B.** The dispersed Golgi in cells described in A was quantified. More than 200 cells were imaged per experiment. Statistical significance was computed by Student’s *t*-test, ***P<0.001. Error bars represent mean ± SEM from three independent experiments. **C&D**. A549 cells stably expressing either control vector or CARP2 without any tag were imaged live with SiaT-RFP marker. Nucleus was stained with DAPI (nucleus). At least 50 cells were imaged per experiment. Statistical significance was computed by Student’s *t*-test, **P<0.001. Error bars represent mean ± SEM from three independent experiments. Scale 10µm. **E**. HEK293T cells stably expressing either control vector or CARP2 stable were transfected with SiaT-RFP. After 24h of transfection the cells were analyzed by FACS and pulse width histograms were represented. Nucleus staining (Hoechst stain) was used as negative control. Statistical significance was computed by Student’s *t*-test, **P<0.001. Data is representative of three independent experiments. **F&G.** HEK293T cells stably expressing CARP2 were transfected with control or CARP2-specific siRNA for 48 hours followed by SiaT-RFP (Golgi marker) live imaging. Nucleus was stained with a Hoechst stain. At least 200 cells were imaged per experiment. Statistical significance was computed by Student’s *t*-test, **P<0.001. Error bars represent mean ± SEM from three independent experiments. Scale 5µm. Western blotting from extracts of these cells were probed with indicated antibodies.

To evaluate whether the effect of CARP2 expression on the Golgi structure is limited to A549 cells or is cell type independent, we also generated HEK 293T cells stably expressing untagged CARP2. Architecture of the Golgi in these cells was evaluated using SiaT-RFP. Whereas the RFP signal was mostly restricted to the perinuclear region in control cells, in CARP2 expressing HEK 293T cells RFP fluorescence appeared as dispersed, like in A549 cells (Figure 2C & D, S1B).

Further, we monitored distribution of the Golgi signal at different intracellular locations using pulse width analysis (PulSA) approach, which measures the width of the fluorescent signal of individual cells in a population by flow cytometry^22–24^. For this analysis we used CARP2 stably expressing HEK 293T cells transfected with SiaT-RFP. Shift in pulse width of SiaT-RFP was observed in CARP2 expressing cells compared to control, confirming that CARP2 promotes Golgi dispersal (Figure 2E).

To ensure the Golgi dispersal observed was indeed because of increased CARP2 protein levels, we reduced the protein level of CARP2 in CARP2-stably expressing HEK293T cells using CARP2-specific siRNA and monitored the Golgi dispersal using SiaT-RFP as a marker. In agreement with a role for CARP2 in maintaining the Golgi structure, transfection with CARP2-specific siRNA, but not control siRNA, restored compact Golgi structure in these cells (Figure 2F & G). CARP2-specific siRNA, unlike control siRNA, was effective in reducing CARP2 protein levels as assessed by western blotting of the cellular extracts from these experiments ( Inset of Figure 2F). These data collectively demonstrate that CARP2 ubiquitin ligase regulates Golgi dynamics.

Since EGF promotes CARP2 upregulation and is reported to be required for cell migration^19^, and EGF-induced cell migration involves remodeling of cell organelles and cytoskeleton^14^, we wondered whether CARP2 expression could also affect, other than the Golgi, the architecture of organelle like mitochondria, ER, and cytoskeleton proteins (Figure S2A-E). For this, A549 cells stably expressing untagged CARP2 were imaged after staining with MitoTracker Red CMXRos (Figure S2B), or after transfecting with an endoplasmic marker (GFP-b5 ER) (Figure S2C), or after immunostaining with a Golgi marker (GM130) (Figure S2A). While no obvious differences were observed in the morphology of either the mitochondria or the ER network in control vs CARP2 expressing cells, visible differences were noted in the compactness of the Golgi apparatus. In these cells the organization of cytoskeleton proteins were also assessed by staining with tubulin antibody (for tubulin) or phalloidin (for actin). No obvious differences in either of these cytoskeleton proteins patterns were noted between control and CARP2 expressing cells (Figure S2D & E). These results suggest that CARP2 specifically regulates the Golgi structure.

### CARP2 ubiquitin ligase activity is essential for the Golgi dispersal

Since, CARP2 is a RING domain protein, which confers ubiquitin ligase function, we were curious to know whether CARP2 ligase activity is required for the Golgi dispersal. For this A549 cells stably expressing CARP2 ligase inactive mutant (CARP2 H333A) were used. Immunostaining of these cells for GM130 showed compact and perinuclear Golgi, as that in vector control cells, suggesting that CARP2 ligase activity is indeed essential for dispersal (Figure 3A & B, S3). In addition to the RING domain, CARP2 contains a FYVE-like motif. This FYVE-like motif is believed to facilitate anchoring of CARP2 on phospholipids with the help of modifications like palmitoylation in the N-terminus. Amino acid cysteine at positions 5 and 6 are the residues predicted to be involved in palmitoylation. Our lab along with other groups showed that mutation of C5,6 to A disrupts the CARP2 localization to intracellular vesicles^25^. We have used this variant of CARP2 to assess the importance of CARP2 endosomal localization for Golgi dispersal. GM130 distribution in C5,6A expressing cells appeared like vector control cells (compact and perinuclear) than CARP2 wild type expressing cells, indicating that CARP2 endosomal localization is required for Golgi dispersal (Figure 3A & B, S3). Like GM130, SiaT-RFP distribution in A549 control, H333A and C5,6A mutant appeared more compact than CARP2 WT expressing cells (Figure 3C & D). These observations indicate that CARP2 ubiquitin ligase activity and endosomal localization play an important role in Golgi structure modulation.

**Figure 3.**
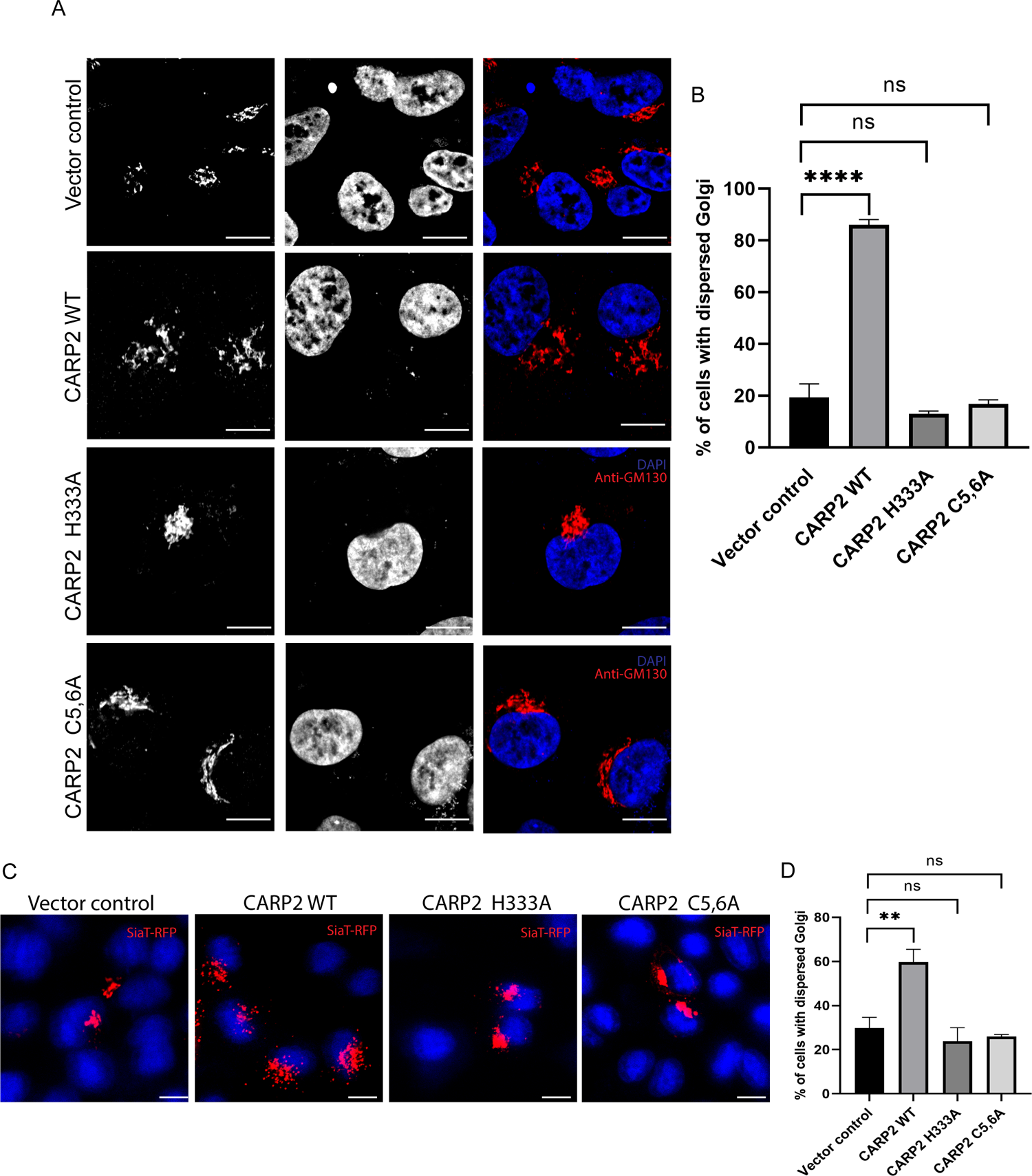
CARP2 ubiquitin ligase activity is required for Golgi dispersal. **A.** A549 cells stably expressing either vector control or CARP2WT or CARP2 H333A or CARP2 C5,6A without any tag were immunostained anti-GM130 antibodies (red). Nucleus is stained with DAPI (blue). Scale 10µm. **B.** The dispersed Golgi in cells described in A was quantified. More than 200 cells were imaged per experiment. Statistical significance was computed by ANOVA, ****P<0.001. Error bars represent mean ± SEM from three independent experiments. **C.** A549 cells stably expressing either vector control or CARP2WT or CARP2 H333A or CARP2 C5,6A without any tag were transfected with SiaT-RFP and imaged live (red). Nucleus is stained with DAPI (blue). Scale 10µm. **D.** The dispersed Golgi in cells described in C was quantified. More than 200 cells were imaged per experiment. Statistical significance was computed by ANOVA, **P<0.001. Error bars represent mean ± SEM from three independent experiments.

### Effect of CARP2 on protein and lipid trafficking

The Golgi is central to the secretory system where packaging, post translational modifications and targeting of lipids and proteins from the endoplasmic reticulum (ER) to the plasma membrane occurs^3^. Having demonstrated that CARP2 expression leads to the Golgi dispersal, we wanted to know whether this dispersal affects the secretion of proteins from the ER. To address the same, we used soluble SS-HRP (SS-HRP oxidase) protein as a surrogate marker. Having a specific signal sequence (SS) that enables Horseradish peroxidase (HRP) to reach the ER, SS-HRP translocate to the ER where the signal sequence is cleaved and the HRP protein is transported outside the plasma membrane through the Golgi bodies. Activity of the HRP secreted into the media is measured using luminol as a substrate^26^. To evaluate the effect of CARP2 expression on the secretion of HRP, HEK293T cells were transfected with CARP2 variants along with SS-HRP-Flag constructs and after 24 hours of transfection the activity of the HRP in the was measured. Increased HRP activity was noted in the media of cells expressing CARP2 WT compared to that of the vector expressing control cells (Figure 4A). Moreover, we could observe dose-dependent enhancement in the HRP activity with an increase in CARP2 expression (Figure 4B). Since CARP2 ubiquitin ligase inactive mutant H333A is unable to affect Golgi integrity, we next measured HRP activity from the media from cells expressing CARP2 H333A mutant (Figure 4C). The intracellular SS-HRP-Flag protein levels in these cells appeared more or less the same, as assessed by immunoblotting with anti-flag antibody (Figure 4A-C, insets). Unlike CARP2 WT, H333A expression did not show any enhanced HRP activity, suggesting that under these experimental conditions, CARP2-induced dispersal of the Golgi results in enhanced secretion of the HRP and this requires the ubiquitin ligase activity. In fact, expression of CARP2 H333A mutant was reported to facilitate accumulation of vesicular structures, which could explain a reduction below the basal level of HRP in these cells^27^.

**Figure 4.**
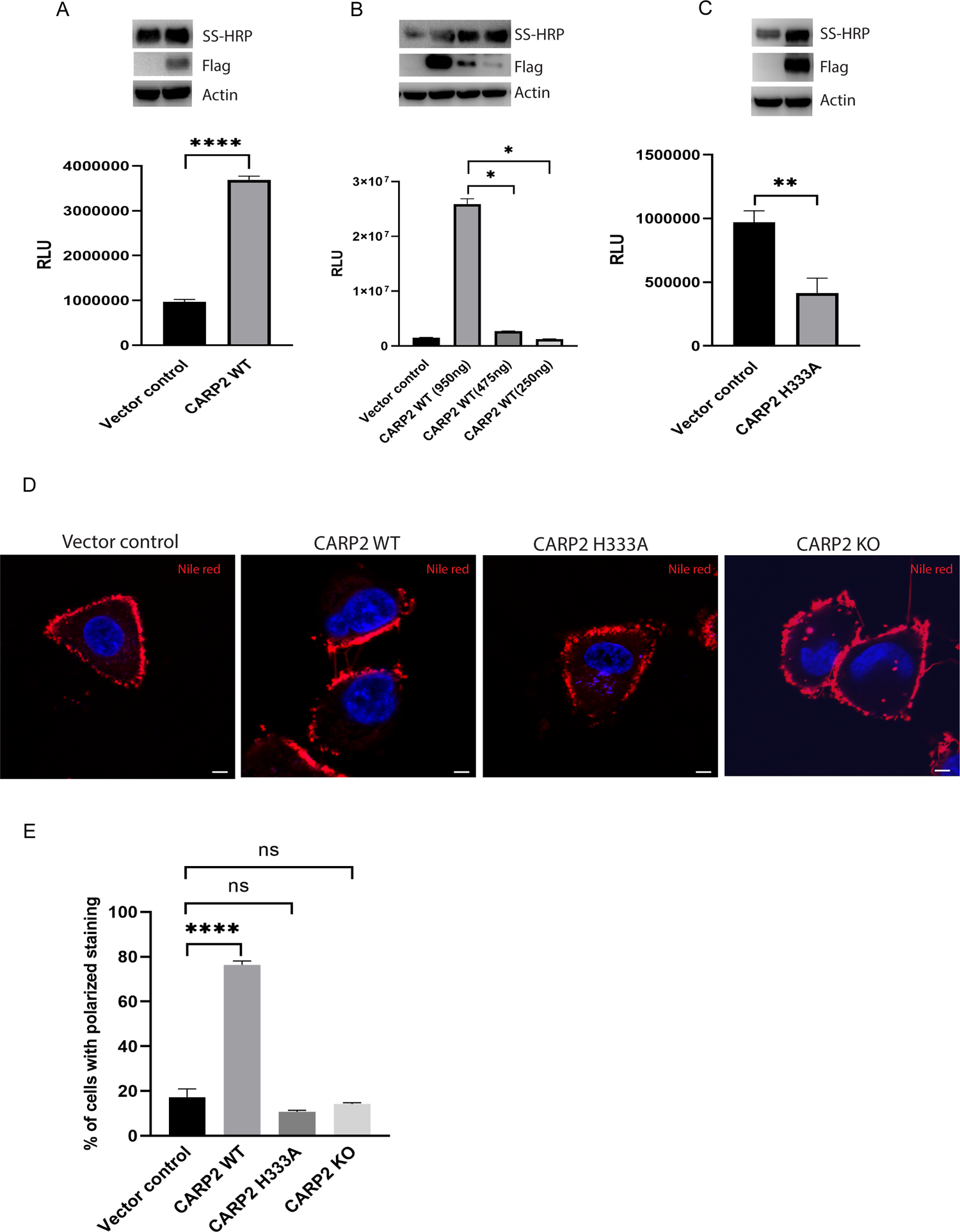
CARP2 affects lipid and protein trafficking. **A.** HRP activity in extracellular media from HEK293T cells transfected with SS-Flag-HRP along with either control vector or Flag CARP2 WT was measured using ECL kit, after 24 hours of transfection. Graph shows the HRP activity from media from cells transfected with different concentrations of Flag CARP2 WT **(B)** and Flag CARP2 H333A **(C)**. Quantification of HRP activity described in A, B, C was computed by ANOVA or Student’s *t*-test. ****,**,*P<0.001. Error bars represent mean ± SEM from three independent experiments. Lysates were subjected to western blotting and probed with indicated antibodies as shown in inset (A-C) **D.** A549 cells stably expressing control vector or CARP2 WT or H333A without tag and CARP2 KO cells were stained with Nile red (red) and nucleus with Hoechst stain, and imaged live. Scale is 5µm. **E.** Quantification of Nile red staining described in D. Statistical analysis was computed by ANOVA, ****P<0.001. Error bars represent mean ± SEM from three independent experiments.

To understand whether CARP2 regulates lipid trafficking or distribution we used the Nile red stain^28^. The Nile red staining of A549 cells stably expressing the control vector or CARP2 H333A, or CARP2 KO revealed staining of intracellular structures appeared as punctate, and present juxtaposed to plasma membrane uniformly. Whereas cells expressing CARP2 WT, staining was more prominently polarized to some areas near the plasma membrane. Nile red stained polarized pattern was quantified and presented (Figure 4D & E). These data suggest the expression of CARP2 WT results in polarized Nile red positive lipid distribution. This polarization requires CARP2 ligase activity.

### Golgin45, a structural protein of Golgi interacts with CARP2

To identify the mechanism by which CARP2 can affect Golgi dynamics, we looked out for CARP2 associated proteins using the Tap-tag system^29^. For this we have expressed CARP2 in 293T cells and the endogenous interaction partners were identified by mass spec analysis We found Golgin45 (also known as BLZF1) as one of the CARP2-associating Golgi structural proteins. Golgin45 is a Golgi tethering molecule with a central coiled coil region that is reported to localize to medial Golgi. Golgin45 is also known to interact with other Golgi membrane adhesion factors like GM130, GRASP55, Syntaxin5 and regulate the Golgi structural dynamics^13, 30^. To confirm whether the Golgin45 indeed is an interacting partner of CARP2, we have performed immunoprecipitation experiments. We transfected 293T cells with Flag-CARP2 and Golgin45-V5-6Xhis constructs and cell lysates were immunoprecipitated with Flag antibody and the precipitates were run on a gel, blotted and probed with anti-V5 antibody. The results showed presence of Golgin45 in the precipitates of CARP2, demonstrating that Golgin45 and CARP2 associate with each other (Figure 5A). Interestingly, the Golgin45 protein levels were consistently observed to be reduced in the lysates of cells expressing CARP2 wildtype, but not H333A mutant (Figure 5B). Given that CARP2 is a ubiquitin ligase, it is likely that CARP2 ubiquitinates Golgin45 leading its elimination. To demonstrate that CARP2 can ubiquitinate Golgin45 in cells, we expressed Golgin45-V5-6xHis and HA-Ub (WT) with or without FLAG-CARP2 WT. The cell lysates were incubated with Ni-NTA beads and the pulldown was evaluated for HA-Ub by probing with anti HA antibody. More HA-ubiquitinated Golgin45 was observed in the presence of CARP2 (Figure 5C). These results indicate that Golgin45 is modified with ubiquitin by CARP2 leading to its degradation.

**Figure 5.**
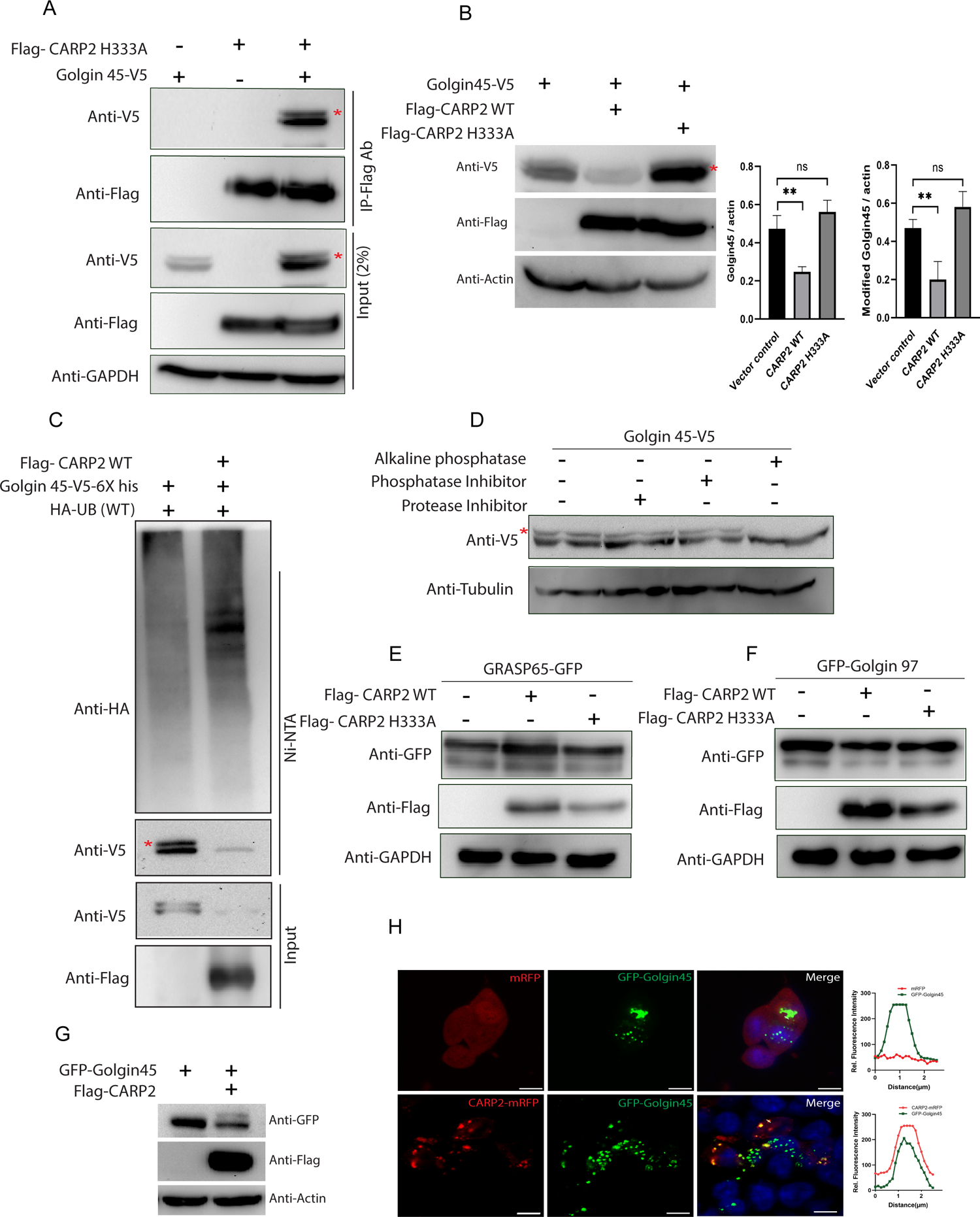
CARP2 interacts and downregulates Golgin45 protein level via ubiquitination. **A.** Cellular extracts from HEK293T cells transfected with indicated constructs were subjected to immunoprecipitation with anti-Flag antibody and the precipitates (IP) along with extracts (input) were probed for Golgin45-V5 with anti-V5 antibody. **B.** Cellular extracts from HEK293T cells transfected with indicated constructs were probed with denoted antibodies. Modified Golgin45 band denoted with * (migrate slower than the unmodified). Quantification of Golgin45 levels after normalizing with GAPDH. Statistical analysis was computed by Student’s *t*-test, *,**P<0.001. Error bars represent mean ± S.D from at least three independent experiments. **C.** Cellular extracts from HEK293T cells transfected with indicated constructs were subjected to Ni-NTA pulldown and probed with denoted antibodies. **D.** Lysates from HEK293T cells transfected with Golgin45-V5, after treatment with alkaline phosphatase for 1 hour, at 37^°^C were probed with anti-V5 and anti-tubulin antibodies. Modified Golgin45 bands were denoted with*. **E&F.** Cellular extracts from HEK293T cells transfected with indicated constructs were probed with denoted antibodies. **G.** Cellular extracts from HEK293T cells transfected with indicated constructs were probed with denoted antibodies. Blot shown is representative from three independent repeats. **H.** HEK293T cells stably expressing either mRFP or CARP2-mRFP were transfected with GFP-Golgin45 or GFP and cells were imaged live. Nucleus (blue) is labelled with Hoechst stain. Scale bar 10µm. Corresponding line profile of colocalized signal is shown.

Interestingly, while probing the cellular extracts with V5 antibody, we noticed the presence of an additional band just above the expected Golgin45 protein band. It has been reported that Golgi structural proteins get phosphorylated to promote Golgi dissassembly^9, 31^. Based on this, we reasoned that the modified Golgin45 band could be phosphorylated. To confirm this Golgin45-V5 lysate was treated with alkaline phosphatase. Interestingly, under treatment conditions, modified Golgin45 is much reduced compared to control lanes (Figure 5D). Thus, we conclude that modified band represents phosphorylated Golgin45. Interestingly, phosphorylated Golgin45 is preferentially reduced in the presence of CARP2. To understand whether CARP2 targeting is specific for Golgin45, protein levels of other Golgi structural proteins GRASP65 (cis Golgi structural protein) and Golgin97 (trans Golgi network structural protein) were assessed in the presence of CARP2 WT and CARP2 H333A mutant. Unlike Golgin45, no obvious difference in the protein levels of GARSP65 and Golgin97 were observed with or without CARP2 (Figure 5E & F)^6, 11^.

For these experiments we used Golgin45 with C-terminus V5-6XHis tag. To avoid any spurious results because of the C-terminus tagging of Golgin45, we have repeated some of the above experiments using N–terminus GFP tagged Golgin45 (GFP-Golgin45). Consistent with results Golgin45-V5, a substantial reduction was also observed in GFP-Golgin45 protein levels when co-expressed with FLAG-CARP2 WT (Figure 5G).

To assess the co-localization between these two proteins, HEK293T cells stably expressing mRFP or CARP2(WT)-mRFP were transfected with GFP-Golgin45. Imaging of cells from these experiments revealed substantial colocalization between GFP-Golgin45 and CARP2-mRFP in structures that appeared to be dispersed Golgi (Figure 5H). We also noticed significant amount of GFP-Golgin45 in the nucleus as reported earlier^32^. These experiments demonstrated that CARP2 specifically targets Golgin45 for ubiquitination leading to its degradation.

### EGF-CARP2-Golgin45 axis regulates the Golgi dispersal

The data thus far demonstrated that: 1. EGF stimulation leads to increased CARP2 protein levels and the Golgi dispersal, 2. Exogenous CARP2 expression results in the Golgi dispersal, 3. EGF induced Golgi dispersal requires CARP2, and 4. CARP2 interacts and ubiquitinates Golgin45 resulting in its degradation. Hence, we hypothesized that endogenous Golgin45 protein levels decrease in response to EGF treatment in CARP2 dependent manner. Towards this end, we analyzed the levels of Golgin45 protein in wildtype and CARP2 KO cells upon EGF stimulation. We found that correlating with an increase in CARP2 protein levels in response to EGF stimulation, a reduction of Gogin45 levels was observed in wildtype cells. However, in CARP2 KO cells Golgin45 remains stable even after three hours of EGF treatment, indicating that EGF stimulation leads to decrease in Golgin45 protein levels and this reduction requires CARP2 (Figure 6A). To further validate the CARP2 mediated Golgi dispersal depends on the reduction of Gogin45 protein. We overexpressed GFP or GFP-Golgin45 in CARP2 stably expressing cells along with SiaT-RFP. Consistent with the idea that CARP2 regulates the Golgi architecture, in GFP expressing cells we found dispersed Siat-RFP. Importantly, increased expression of GFP-Golgin45 resulted in more compact SiaT-RFP staining in the perinuclear region (Figure 6B). These results indicate that CARP2 regulates the Golgi structure by targeting Gogin45. Overall, these findings reveal hitherto unknown CARP2-Golgin45-Golgi axis in EGF stimulated cell migration pathway.

**Figure 6.**
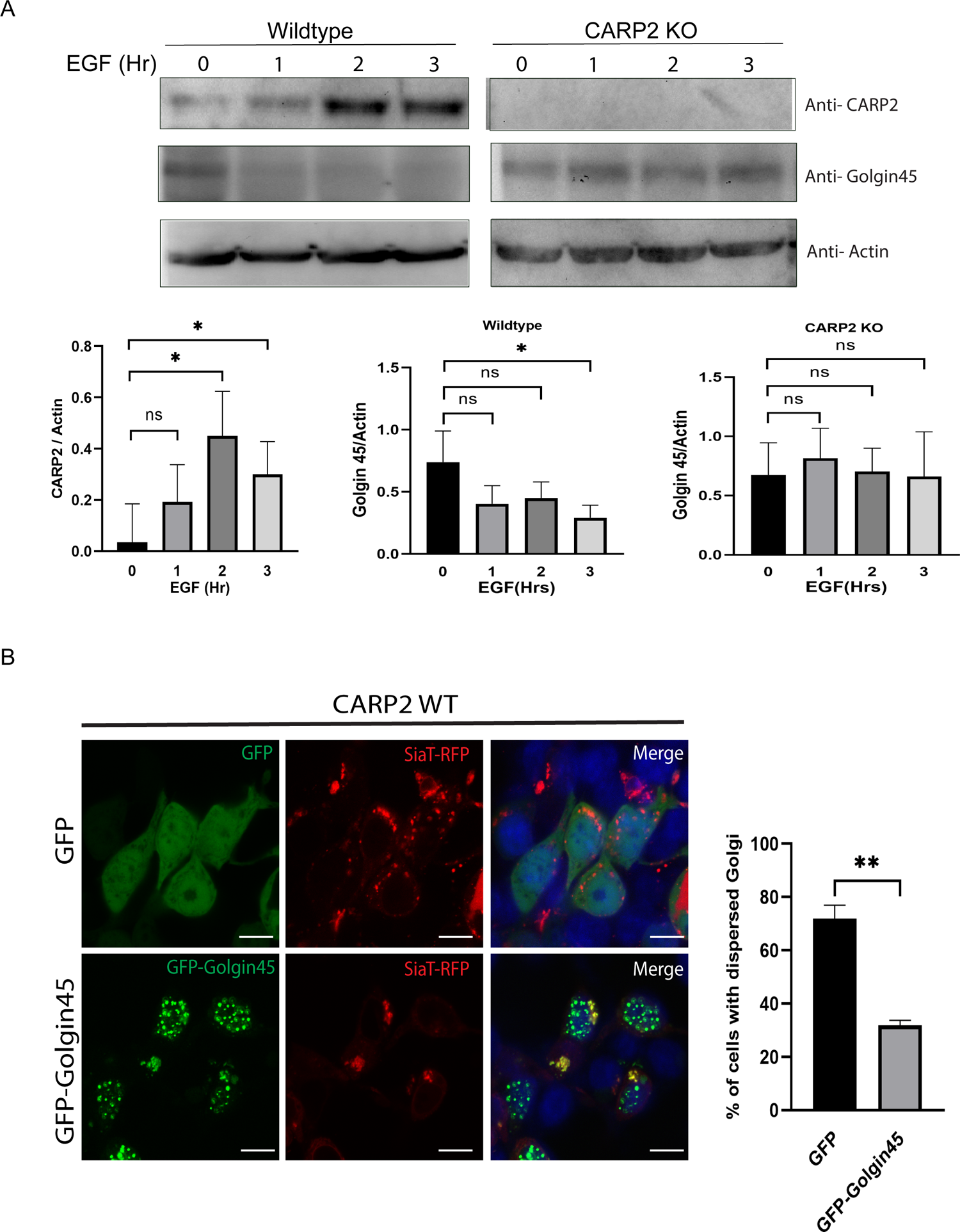
EGF-CARP2-Golgin45 axis regulates the Golgi dispersal. **A.** A549 wildtype or CARP2 KO cells were treated with EGF for indicated time periods and extracts were probed with indicated antibodies (top panel) and quantification of CARP2, and Golgin45 levels after normalization with actin shown (bottom panel). Statistical analysis was computed by ANOVA, *P<0.001. Error bars represent mean ± S.D from three independent experiments. **B.** HEK293T cells stably expressing CARP2 without any tag were co-transfected either with GFP and SiaT-RFP or GFP-Golgin45 and SiaT-RFP followed by live imaging. Nucleus (blue) is labelled with Hoechst stain. Scale bar 10µm. The dispersed Golgi in cells was quantified. Statistical analysis was computed by Student’s t test, ***,*P<0.001. Error bars represent mean ± SEM from three independent experiments. At least 100 cells were imaged for each biological replicate.

## Discussion

Delicate balance between vesicle fission and fusion events governs the Golgi structural dynamics and function^33^. Converging evidence indicates that Golgi structural remodeling occurs in several physiological events including cell division and migration. In dividing cells, the Golgi ribbon undergoes disassembly by sequential unlinking and unstacking followed by reassembly after cytokinesis, which is essential for segregation of Golgi among daughter cells^3^. This is ensured with the help of cell cycle dependent kinases, phosphatases, and ubiquitin ligases. In migrating cells, unlinked Golgi apparatus redistribute to perinuclear sites towards the leading edge of the cell^4, 8^. Dysregulation of the Golgi dynamics and fragmentation is reported in several pathological conditions like cancer and neurological disorders^34^. Here, we presented evidence that increased levels of CARP2 protein leads to dispersal of compact Golgi into mini-stacks, as observed by GM130 (cis-Golgi marker) and SiaT-RFP (trans Golgi marker) distribution. Depletion of CARP2 using CARP2 specific siRNA reversed the dispersed Golgi phenotype observed in CARP2 stably expressing cells, demonstrating the specificity of CARP2 effect on Golgi structural integrity. Further, CARP2 mediated Golgi dispersal phenotype is not restricted to A549 cells but similar distribution was observed upon exogenous expression of CARP2 in other cell types like HEK293T cells and N2A (data not shown). Moreover, CARP2 ubiquitin ligase activity is essential for Golgi dispersal because CARP2 ligase inactive mutant is unable to cause Golgi dispersal. Given that CARP2 expression is reportedly high in brain and gonadal tissues^35^, it would be interesting to investigate whether differential expression of CARP2 in a given cell or tissue type could be linked to cell or tissue specific variations in Golgi morphologies.

While deciphering the molecular mechanism underlying CARP2 mediated Golgi dispersal, we identified Golgin45 as an interacting partner by Tandem affinity purification tag approach. We demonstrated that CARP2 interacts with Golgin45, and ubiquitinates and downregulates its protein level in ubiquitin ligase-dependent manner. Interestingly, we also found a modified Golgin45 variant that is sensitive to alkaline phosphatase and susceptible to CARP2-mediated degradation. These observations are consistent with earlier reports that suggested downregulation of Golgin45 by siRNA leads to Golgi dispersal. Depletion of Golgin45 results in an increase in cisternae luminal width, and a reduction in the average number of cisternae per stack as observed by electron microscopy in HeLa cells^30, 36^. Importantly, the phenotype of the dispersal of Golgi in CARP2 WT stably expressing cells could partly be reversed upon Golgin45 overexpression indicating signaling between Golgin45 and CARP2 determines the Golgi architecture. These results collectively suggest that CARP2 mediated Golgi dispersal is a consequence of Golgin45 downregulation. While CARP2 controls the Golgi architecture by targeting Golgin45, we envisage that CARP2 modulates the Golgi apparatus by regulating several other Golgi structural proteins. The CARP2 TAP-tag screening identified ACBD3 and COG proteins, molecules that are known to control the Golgi dynamics. Though not explored in the context of the present investigation, it would be interesting to study the effect of CARP2 on ACBD3 and COG proteins with reference to the Golgi architecture. Another important aspect that warrants further investigations is Golgin45 phosphorylation. Studies based on high throughput proteomics reported that phosphorylation of Golgi structural proteins including Golgin45 occurs during mitosis correlating with Golgi disassembly^37–39^. We are not aware of the details of kinases that modify Golgin45 or the effect of this modification on its function. It is logical to think that rapid modification and turnover of these structural proteins could facilitate quick rearrangement of the Golgi structure under physiological conditions. It would also be interesting to explore the role of CARP2 during Golgi restructuring in mitotic cells.

The ER-Golgi system plays a central role in modification of proteins with moieties like glycosylation, acetylation and their sorting to different destinations through vesicular trafficking and secretion. This process is complex and is regulated at multiple levels depending on the biochemical nature of the cargo such as proteins or lipids and their size, solubility and topology. Some of the structural proteins like golgins also facilitate cargo transport in addition to giving structural integrity to the Golgi. For example Golgin97 promotes transportation of E-cadherin while Golgin245 facilitates that of TNF-alpha suggesting the existence of diverse transportation routes^40, 41^.

Maintenance of the integrity of the Golgi is required for cellular homeostasis as the dispersal of the Golgi was reported under pathological conditions like neurodegeneration and cancer^34^. Loss of the Golgi ribbon structure alters intracellular trafficking of both proteins and lipids with reports suggesting both increase or decrease in the trafficking. Increased secretion of tau and amyloid beta proteins observed in case of Alzheimer’s diseases associated with dispersed Golgi. Similarly, depletion of GRASP proteins is associated with the Golgi dispersal and enhanced CD8 and cathepsin D trafficking^42, 43^. On the contrary, loss of GMAP-210 (cis-golgin) or Giantin (medial-golgin) in mouse which is associated with altered Golgi also reported to decrease in the secretion of extracellular components^31, 44^. Golgin45 has also been reported to control the trafficking of small cargoes^45^.

Consistent with these reports, CARP2 expressing cells showed along with dispersed Golgi an increased secretion of SS-HRP cargo. One interesting observation, though inexplicable at this stage, is change in near to plasma membrane distribution of Nile red staining in CARP2 expressing A549 cells. Nile red is known to stain lipid droplets, particularly hydrophobic lipids like cholesterol^28^. The plasma membrane is enriched with cholesterol where it constitutes to nearly 25% of the total lipid content. The distribution of lipids/cholesterol affects the biophysical properties like fluidity and the permeability of the membrane. Polarized distribution of lipids in cell membranes is believed to be required for specialized functions, like signalling, membrane trafficking, and cell adhesion^46–49^. More investigations are required to understand the role of CARP2 in the lipid distribution, as exemplified by Nile red distribution, and network of endosomal-Golgi pathways involved in the process.

Importantly, we also demonstrated that the CARP2-Golgin45-Golgi axis is physiologically relevant to EGF-induced cell migration. Earlier we reported an increase in the transcriptional upregulation of CARP2 in EGF treated cells^19^. Interestingly, EGF stimulation is also associated with the Golgi fragmentation. In this paper we presented evidence that in EGF stimulated cells, increase in CARP2 protein level correlated with decrease in endogenous Golgin45 protein. This reduction in Golgin45 protein requires CARP2, as no downregulation was noted in CARP2 KO cells. Moreover, we demonstrated that absence of CARP2 reduces EGF-dependent SiaT-RFP dispersal. We also found that exogenous expression of GFP-Golgin45 attenuated CARP2-induced Golgi dispersal. Furthermore, increased transcript levels of CARP2 are also reported in bleomycin induced pulmonary fibrosis^20^. It is interesting to note that pulmonary fibrosis is manifested by large secretion of extracellular matrix components. Our findings raise potential involvement of CARP2 in the regulation of diverse physiological processes like cell migration and division, protein trafficking. It would be interesting to investigate CARP2-mediated regulation of the Golgi dynamics, secretion and polarization under pathological conditions like cancer. With recent evidence indicating that dispersed Golgi facilitated the development of drug resistant tumors and increased CARP2 mRNA expression was attributed to resistance to chemotherapy induced cell death in primary tumor cell lines^50^, it would be important to investigate role of CARP2-mediated Golgi dispersal in the development of resistance to multi drug therapy by resistant tumor cells. Our findings support an important role for CARP2 in the Golgi dynamics and reveal complex crosstalk between endo-lysosomal systems and the Golgi apparatus.

## Acknowledgments

We thank CSIR for the financial support to RS. This project was supported by IISERTVM intramural grant, Department of Science and Technology-Science and Engineering Research Board, India (DST-SERB) grant (EMR/2016/008048), and Department of Biotechnology grant (BT/PR21325/BRB/10/1554/2016), and STAR grant STARS/APR2019/BS/708/FS awarded to SMS. We are thankful to Rishith Ravindran, Dr. Anoop Kumar, Nikhil Dev for sharing the valuable reagents and Dr. Jishy Varghese for sharing the Nile red stain. We would like to acknowledge Sarika Mohan and Irene Mariam Roy for their technical assistance with flow cytometry.

## Author contributions

Conceptualization: RS & SMS; supervision: SMS; experiment designing: RS & SMS; experiments performed: RS; data analysis: RS & SMS; manuscript preparation: RS & SMS; project administration: SMS

## Data availability

All the data generated and analyzed are included in this manuscript.

## Consent for Publication

Authors agreed to publish this manuscript.

## Ethics Approval

HEK293T and A549 cells used in this study were obtained from ATCC and approved by the institute biosafety committee.

## Supplementary

**Figure 1.**

**A.** A549 cells stably expressing either vector control or CARP2 WT without any tag were cells immunostained with anti-CARP2 antibody (green). Scale bar is 10µm. Protein levels are shown in respective insets by immunoblotting. **B.** HEK293T cells expressing control vector or CARP2 WT were transfected with SiaT-RFP (Golgi marker) and nucleus (blue) is stained by Hoechst stain. Scale 10µm.

**Figure 2.**

**A.** A549 cells expressing control vector or CARP2 WT were immunostained with anti-GM130 (for Golgi) and DAPI stained the nucleus. **B.** Cells described in A were stained with MitoTracker Red CMXRos (for mitochondria) and Hoechst stain (nucleus) and imaged live. **C.** Cells described in A were transfected with GFP-b5 ER (for endoplasmic reticulum) and cells were imaged live. Nucleus was stained with a Hoechst stain. **D.** Cells described in A were immunostained with anti-tubulin antibody (for cytoskeleton) and the nucleus with DAPI. **E.** Cells described in A were stained with phalloidin (actin marker) and Hoechst stain (nucleus) and imaged live. Scale 10µm. These are representative images from three independent experiments.

**Figure 3.**

A549 cells stably expressing either vector control or CARP2WT or CARP2 H333A or CARP2 C5,6A without any tag or CARP2 KO cells were immunostained with GM130 antibody (red). Nucleus was stained with DAPI (nucleus). Scale 10µm. Images are representative from five independent experiments.

## Materials and Methods

### Cell culture and EGF Stimulation

Cell lines used in this study were obtained from ATCC. Cells were grown in Dulbecco’s modified Eagle’s medium (DMEM with 4.5 g/l glucose, L-glutamine, 3.7 g/l sodium bicarbonate and sodium pyruvate) (SH30243.01, Hyclone GE) supplemented with 10% FBS (10270-106, Gibco-Thermo Fisher Scientific) and 1% penicillin-streptomycin (A001A, HiMedia). Cells were kept in at 37°C supplied with humid 5% CO2. Periodically, cells were tested for contamination. Cells were plated 24 hours prior to transfection and Lipofectamine 3000 was used to transfect the plasmid as per the manufacture’s instruction (L3000-015, Invitrogen). CARP2 variants stable cell lines (A549, HEK293T) were generated using retroviral method followed by selection with puromycin(1ug/ml) as described earlier. CARP2 knock cells were generated in our lab using CRISPR^23^. For EGF stimulation experiment, A549 cells were serum starved for 16 hours followed by addition of EGF (200 ng/ml) for a given time period.

### Plasmids and Reagents

Human CARP2 cDNA (NCBI reference sequence: NM_001017368.2:141- 1232) was cloned in pFlagCMV2 vector and pMYs-IP. Golgin45-V5 (Harvard plasmid repository), GFP-Golgin97 (Chia-Jung Yu, Chang Gung University, Taiwan), GRASP65-GFP (Yanzhuang Wang, University of Michigan, USA) and SiaT-RFP (Jack Rohrer, ZHAW, Switzerland), GFP-Golgin45 (Intaek Lee, ShanghaiTech University, China) and SS-HRP-Flag (Frederic Bard, IMCB A*STAR, Singapore) constructs received as generous gift.

Antibodies GM130 (G7295, Sigma), GFP (632375, Mouse Living Colours Clonetek), Tubulin (T6199, Sigma-Aldrich), Actin (4967S, CST), Flag (F3165-1MG, Sigma), V5 (R96125, Invitrogen), BLZF1 antibody (MA5- 27126, Invitrogen), Anti mouse HRP (712-035-150, Jackson Immuno Research), Anti-rabbit HRP (65-6120, Invitrogen) were purchased. Epidermal Growth factor (E9644, Sigma). Polyclonal CARP2 antibody raised in Rabbit (generated in our laboratory)^25^.

### RNA Interference

To downregulate the CARP2 levels, cells were transfected with 1µM of either control or CARP2 specific siRNA- sequence is − 5’ UGACAUCUCUACCGAAAUG 3’ using RNAimax (13778-075, Thermo Fisher Scientific) as per manufacturer’s instructions.

### Immunoblotting

After 30 hours of transfection cells were lysed in lysis buffer (20mM Tris-cl (pH 7.5), 50mM NaCl, 1mM EGTA, 0.5% Triton X-100, 1X-protease inhibitor cocktail, 1X-phosphatase inhibitor 2 and 3) for 30 mins on ice. Followed by spin at 13,500×g for 10 min and supernatant was collected, samples were prepared using Laemmli buffer and boiled at 95°C for 10 minutes. Samples were separated by SDS-PAGE and later transferred to the PVDF membrane. Immunoblotting was performed with specific antibodies as mentioned in the figures and detection was done with chemiluminescent HRP substrate. (WBKLS0500, Merck). For immunoprecipitation, lysates from transfected cells were prepared as mentioned above, and pulled down with indicated antibodies as shown in figures.

### Immunostaining and Imaging

Cells were grown on coverslips and briefly washed with PBS, followed by fixation with 4% paraformaldehyde for 20 minutes at room temperature. Washed thrice with PBS before permeabilization with 0.1% triton-X100 for 10 minutes. Then washed again, blocked with 3% bovine serum albumin prepared in PBS for 1 hour, followed by incubation with primary antibody and washing. Secondary antibodies conjugated with fluorophore against primary antibody was incubated for 1 hour and washed with PBS and nucleus was stained with DAPI. Then the coverslip was mounted on a glass slide with antifade mounting media.

Next day images were acquired using 63X oil immersion objective of Leica TCS 508 SP5 II laser scanning confocal microscope. For live cell imaging, cells were grown on glass bottom dish (MatTek 254 Corporation, P35G-1.0- 20-C) followed by imaging using the same microscope mentioned earlier under live cell imaging settings. Here, Hoechst stain used to label nucleus. Golgi dispersal quantification was carried out manually with criteria like if signals are scattered dots in perinuclear region or not connected, or isolated dots considered as dispersed Golgi, otherwise intact.

For Nile red staining, cells were plated 24 hours before live imaging in a glass bottom dish. Then, stained with Nile red dye (to stain lipid) for 10 mins and briefly rinsed with PBS. Nucleus was stained by Hoechst dye.

### SS-HRP Secretion Assay

HEK293T cells co-transfected with SS-HRP-Flag with either empty vector or Flag-CARP2WT or Flag-CARP2 H333A plasmids respectively. After 24 hours post transfection conditioned media was used to measure the secreted HRP activity^26^.

### Pulse width analysis

Cells were transfected with plasmids in mentioned cell lines (see legends), after 24 hours cells were collected with PBS supplemented with 5mM EDTA and analyzed at medium flow rate in flow cytometer FACS Aria III (BDBiosciences, San Jose, CA) and FlowJo software was used for analysis. Flow cytometer was equipped with lasers 405, 488 and 561nm. Forward scattered threshold of 5000 was used to collect more than 10,000 events. For each channel pulse height, width, and area parameters were recorded and analyzed the pulse shape as described earlier^22^.

### Statistical Analysis

Experimental data is represented as mean ± s.d or sem. Student’s t test or ANOVA test was performed to obtain p-value, with α=0.05 and significance levels demonstrated as ns, not significant, *P<0.05, **P<0.01, ** P<0.001 and ****P<0.0001. Graph pad (version 8.2.0) software was used. All the experiments were repeated at least three times independently.

