## Supplementary figures and images for "CARP2 regulates the Golgi dynamics upon EGF stimulation"

### Supplementary Figure 1

Supplementary Figure 1

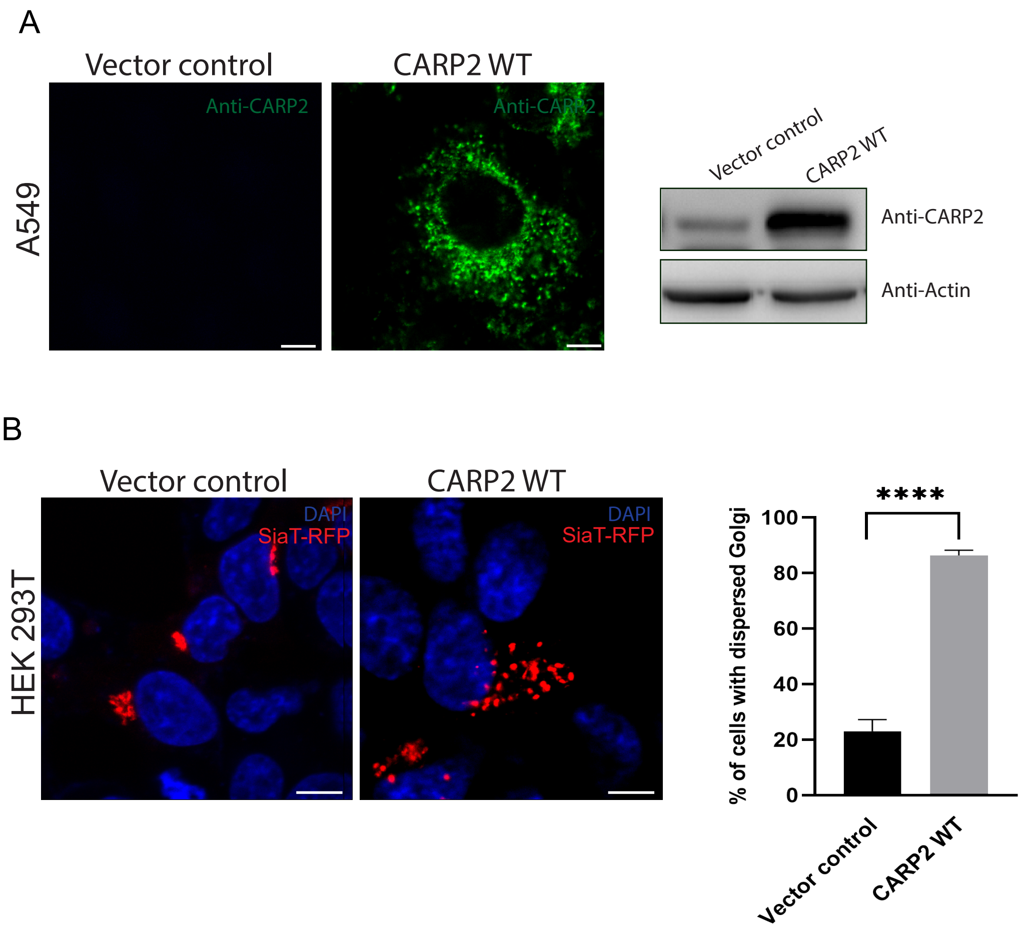

### Supplementary Figure 2

Supplementary Figure 2

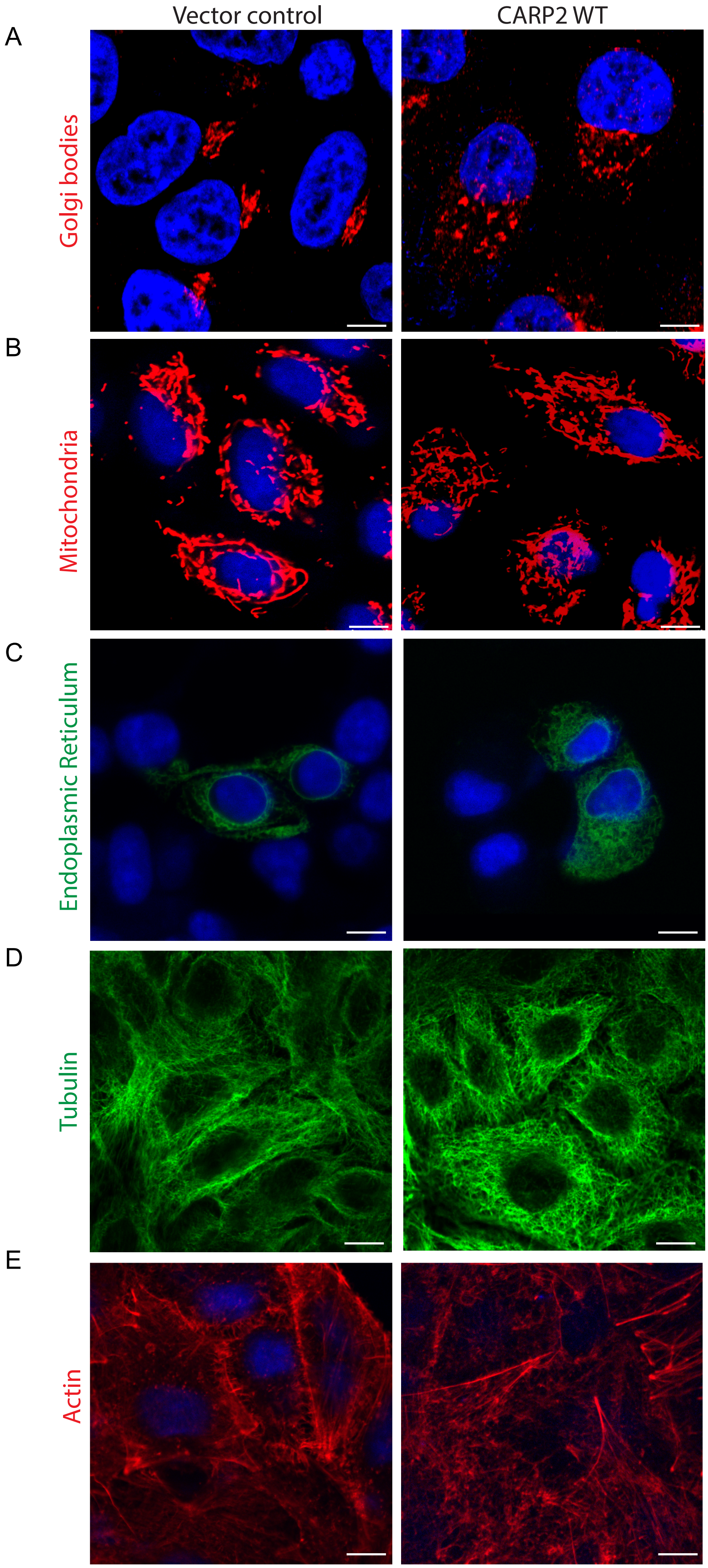

### Supplementary Figure 3

Supplementary Figure 3

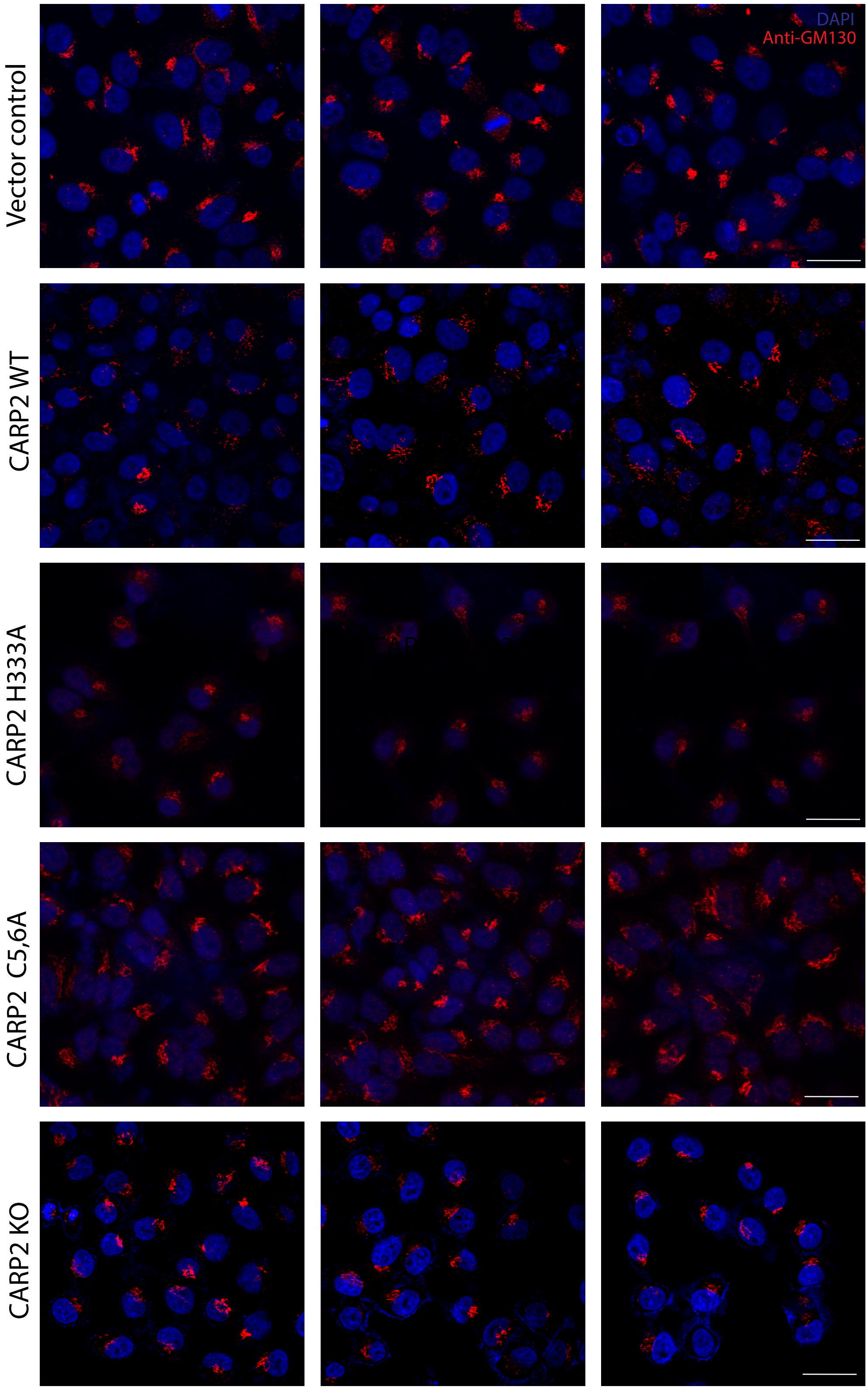
